# Activity flow over resting-state networks shapes cognitive task activations

**DOI:** 10.1101/055194

**Authors:** Michael W. Cole, Takuya Ito, Danielle S. Bassett, Douglas H. Schultz

## Abstract

Resting-state functional connectivity (FC) has helped reveal the intrinsic network organization of the human brain, yet its relevance to cognitive task activations has been unclear. Uncertainty remains despite evidence that resting-state FC patterns are highly similar to cognitive task activation patterns. Identifying the distributed processes that shape localized cognitive task activations may help reveal why resting-state FC is so strongly related to cognitive task activations. We found that estimating task-evoked activity flow (the spread of activation amplitudes) over resting-state FC networks allows prediction of cognitive task activations in a large-scale neural network model. Applying this insight to empirical functional MRI data, we found that cognitive task activations can be predicted in held-out brain regions (and held-out individuals via estimated activity flow over resting-state FC networks. This suggests that task-evoked activity flow over intrinsic networks is a large-scale mechanism explaining the relevance of resting-state FC to cognitive task activations.

---

The neural basis of cognition has been primarily investigated in terms of task-evoked activation level changes. Over the past decade a separate focus on spontaneous (non-task-evoked) activity has challenged cognitive neuroscientists’ focus on task-evoked activations^1 –3^. Due to the lack of experimental control of spontaneous brain activity there has been a strong emphasis on discovering correlations (rather than activation level changes) among activity time series-an approach termed resting-state functional connectivity (FC). Thus, the theoretical framework and methodological approaches associated with cognitive task activations and resting-state FC are highly distinct, leading to a bifurcation in investigations of brain function.

Notably, this bifurcation mirrors the classic “localized” versus “distributed” neural processing debate^4  –7^, such that the relationship between localized cognitive task activations and distributed FC is also relevant to this broader theoretical divide in neuroscience. Here we sought to identify where the human brain lies with respect to these two extremes. We focused in particular on the role of intrinsic functional networks (as estimated by resting-state FC) in distributed processing. There is evidence that resting-state FC patterns are similar to cognitive task activation patterns^8, 9^, but we sought to quantify this relationship using a large-scale mechanistic account that might explain why this relationship exists. Critically, we recently found that the FC architectures across a variety of tasks were highly similar(80% shared variance) to the resting-state FC architecture^10^. This suggests that the network architecture identified using resting-state FCis present during taskperformance, and could plausibly reflect the routes by which activity flows during cognitive task performance. However, it remains unclear whether andhow these FC patternsrelate to cognitive task activation amplitudes-such as task-evoked blood oxygen level dependent (BOLD) functional MRI (fMRI) signal increases-and therefore how they relate to cognition.

We sought to answer these questions by testing whether estimated activity flow over resting-state FC networks can accurately predict held-out cognitive task activations. Activity flow (often termed “information flow”) is the spreading of activation amplitudes between brain locations, such as task-evoked activations spreading from visual cortex to motor cortex in a visual-motor task. Decades of findings in local circuits and simulations have suggested that connectivity and activations are strongly interrelated neurophysiological variables, with activity/information flow as a key linking variable^11 –13^. However, little is known about how FC and cognitive task activations relate at the large-scale network level,e.g., as measured with fMRI. Beginning to fill this gap, several recent studies used abstract statistical models to predict cognitive task activations based on individual differences in large-scale connectivity^9,14, 15^. We sought to build on these findings to identify why these predictions were possible. This involved testing the plausibility of a (large-scale) mechanistic relationship between connectivity and cognitive task activations in terms of the concept of activity flow.

Local circuit-level studies have suggested that task-evoked activation at a given location is primarily determined by activity flow from other neurons^16, 17^. Activityflow is carried (via axons) by action potentials modulated by synaptic strengths. Thus, activity flow is a mechanism that emerges from several more basic mechanisms (action potentials, synaptic strengths, etc.). One can conceptualize activity flow as relating activations (action potentials and associated local field potentials) and functional pathways (their tendency to influence one another via synaptic strengths). Applied to large-scale measures of the human brain, we hypothesized that aggregate activation amplitudes (e.g., as measured by BOLD fMRI signal) flow among brain regions via functional pathways (possibly reflecting aggregate synaptic strengths) described by FC. Thus, we conceptualized activity flow as a linking variable between large-scale FC and cognitive task activations that could be used to demonstrate (and quantify) the functional relevance of these two measures to one another.

We tested the plausibility of this hypothesis by constructing activity flow mappings using FC and task activations. This involved predicting the cognitive task activation level at one location based on the FC-weighted sums of the activations at other locations (**Fig. 1A**). We then repeated this process separately for each brain region-akin to cross-validation from machine learning^18 –20^. This resulted in a whole-brain activation pattern prediction, which could be compared with a given task’s actual fMRI activation pattern. A successful prediction (i.e., high correspondence between predicted and actual activation patterns) would indicate the plausibility of resting-state FC pathways in shaping the empirically observed activation pattern. Further, successful prediction across a variety of tasks and subjects would indicate the generalplausibilityof the activity flow framework at the large-scale network level-suggesting resting-state FC is relevant to cognitive task activations due to its role in shaping task-evoked activity flow among brain regions.

There are several reasons why activity flow-based prediction of cognitive task activations is not guaranteed to work. For instance, cognitive task activations may be largely shaped by task-evoked network reconfigurations^17, 21, 22^, making prediction of cognitive task activations by resting-state FC ineffective. Additionally, localized processing independent of other brain regions could be a major driver of cognitive task activations in any given brain region, such that activity flow is largely irrelevant to localized cognitive task activations. Indeed,many cognitive task activations have been interpreted under this assumption^7^, such as task-evoked activations within dorsolateral prefrontal cortex during working memory maintenance^23^. Even with strong evidence that activity flow shapes activations at the local circuit level^11 –13^, this is not guaranteed at the large-scale network level since local processing (e.g., within-region activity flow) is likely to be at least partially independent of the large-scale activity flow into a region.

The activity flow mapping approach is based on the local circuit-level findings described above. However, like recent models of activity spreading dynamics^24, 25^, it is not meant to be a realistic simulation of neuronal dynamics but rather a tool for quantifying (and making inferences about) brain activity relationships. We see the present study as a precursor to more complex approaches that incorporate biophysical models of neuronal communication^26, 27^ to improve activation pattern predictions further.Here we sought to make as few assumptions as possible regarding the biophysical basis of the FC-activation relationship by using the simplest activity flow mapping approach possible-the FC-weighted sum of activations. This allowed us to make straightforward inferences regarding the relationship between FC and cognitive task activations, which future work can refine using more elaborate models of neuronal communication.

We began by validating the activity flow mapping procedure with a simple computational model of large-scale neural interactions. We then applied the activity flow mapping approach to empirical fMRI data acquired as healthy adult human subjects (N=100) rested and performed a variety of tasks. We used activity flow mapping to test our primary hypothesis: That cognitive task activations can be predicted in held-out brainregions (and held-out individuals) via estimated activity flow over resting-state FC networks. This would suggest that task-evoked activity flow over intrinsic networks (i.e., the spread of activation amplitudes between regions) acts as a large-scale mechanism explaining the relevance of resting-state FC to cognitive task activations.

**Figure 1.**
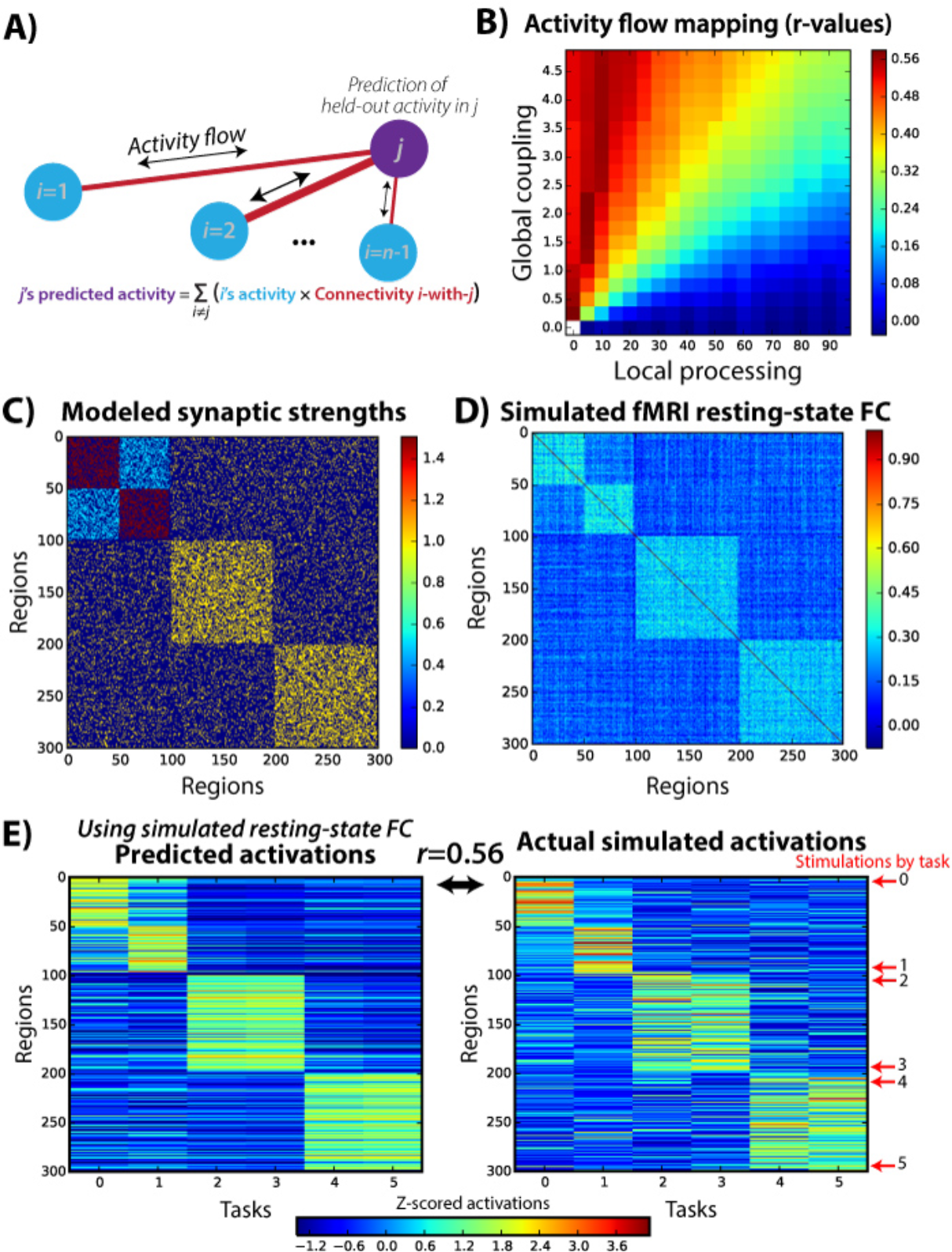
**Activity flow mapping over resting-state FC networks allows prediction of held-out task activations**. **A**) We developed a prediction-based approach that links resting-state FC to task activations to assess the relevance of FC to cognitive task activations. The prediction of a single region’s activation amplitude for a single task is depicted. Importantly, the to-be-predicted region’s activation amplitude is held out from the prediction calculation. **B**) This approach was validated using a simple computational model of large-scale neural activity. Whole-brain predicted-to-actual Pearson correlations (r-values) for distinct model parameters are shown. The success of activity flow mapping depended on the relative degree of local versus distributed processing. This demonstrates that the success of activity flow mapping with empirical fMRI data would be non-trivial. **C**) Three structural connectivity graph communities (blocks along diagonal) were created, with the first split into two communities via synaptic strength modifications. **D**) Resting-state FC (Pearson correlation) was computed based on simulated timeseries using the computational model, revealing a strong correspondence with the underlying synaptic strengths. Both the global coupling and local processing parameters were set to 1.0 for this example. *E*) Simulated task-evoked activations were produced by stimulating groups of 5 nearby units in 6 separate “tasks”. The activity flow mapping procedure produced above-chance recovery (mean across-task Pearson correlation *r*=0.56, t(298)=11.7, *p*<0.00001), of the actual activations using the resting-state FC matrix shown in panel D.

## RESULTS

### Computational validation and identification of factors contributing to cognitive task activations

Previous research has shown that there is a statistical relationship between resting-state FC and cognitive task activations^8, 9^, but not why this relationship exists. We recently found that resting-state FC patterns are present during cognitive task performance (80% shared variance in whole-brain FC patterns between rest and task)^10^. This suggests that resting-state FC might describe activity flow among brain regions even during task performance. Here we tested this possibility in the context of task-evoked activation amplitudes, using activity flow among brain regions as a linking variable between resting-state FC and task-evoked activations. This involved modeling activity flow as task activation amplitudes (standard fMRI general linear model estimates) multiplied by FC strengths (standard Pearson correlations) between brainregions (**Fig. 1A**). Standard measures were used to maximally relate to the existing resting-state FC and cognitive task activation literatures. We hypothesized that this would allow us to predict cognitive task activations in held-out brain regions based on resting-state FC patterns.

We began by validating this activity flow mapping procedure with a simple computational model of large-scale neural interactions. The model was kept simple to reduce the number of assumptions regarding underlying biophysical detail (see Methods). Interactions among 300 brain regions were simulated along with task-evoked activations. Knowing the ground truth connectivity and activations in the model allowed us to validate the activity flow mapping procedure.

We constructed the model to have three structural network communities, with the first community split into two “functional” communities via modulation of synaptic strengths (**Fig. 1C**). This was of particular interest here given the potential for resting-state FC fMRI (unlike, e.g., diffusion-weighted MRI) to detect the aggregate effects of synaptic strengths that are known to modify activity flow over structural (axonal) connections in local circuits^11^. We then ran the model with spontaneous activity (Gaussian random values) in each unit while simulating fMRI data collection (see Methods). We then computed Pearson correlations among all of the time series to produce simulated resting-state FC data (**Fig. 1D**).

We next simulated task-evoked activations by injecting stimulation (2 simulated minutes of stimulation in 3 blocks) into 5 neighboring regions at a time. Six “tasks” were simulated by changing the stimulated regions (see **Fig. 1E**). We simulated fMRI data collection as with the “rest” data, using a standard fMRI general linear model to obtain activation amplitude estimates for each simulated region (**Fig. 1E**). We then ran the activity flow mapping algorithm to assess the ability to use resting-state FC to predict task activations in held-out regions.We found that activity flow mapping was successful in recovering the original task-evoked activation pattern (*r*=0.56, ρ<0.00001; Spearman’s rank correlation rho=0.51, p<0.00001).

To ensure robustness of this result we repeated the entire simulation procedure 4000 times (see Methods). Over these iterations we varied a global coupling parameter to assess the role of aggregate synaptic strengths, along with varying a local processing parameter to assess the role of non-distributed (local) activity. Global coupling was defined as a constant that linearly scaled all synaptic strengths, while local processing was defined as a constant that linearly scaled all self (recurrent) connection strengths. We found that activity flow mapping worked to the extent that global synaptic connectivity was high and local processing was low (**Fig. 1B**). The sensitivity of these results to the local-distributed processing relationship suggested that empirical assessment of activity flow mapping with real fMRI data would be nontrivial, in the sense that it would only be predictive if the signals displayed certain properties. Further,these results suggest that activity flow mapping could provide evidence regarding the relative distributed versus localized processing that occurs in the human brain during cognitive task performance.

### Activity flow mapping with empirical fMRI data

We next applied the activity flow mapping approach to empirical fMRI data, testing the hypothesis that cognitive task activations can be predicted in held-out brain regions via estimated activity flow over resting-state FC networks. This involving applying activity flow mapping to a Human Connectome Project dataset involving rest and 7 highly distinct tasks^28^. The predicted activation pattern matrix that was highly similar to the actual activation pattern matrix (**Fig. 2B**): average *r*=0.54, t(99)=49.95, ρ<0.00001. The r-values were similar for each of the seven tasks individually: *r*=0.56 (emotional), *r*=0.54 (gambling), *r*=0.55 (language), *r*=0.53 (motor), *r*=0.56 (relational), *r*=0.54 (social), *r*=0.54 (N-back).These correlations were higher when computed after averaging the FC and activation patterns across subjects (average *r*=0.60), likely due to an improved signal-to-noise ratio from aggregating more data prior to the comparisons. Note that this average-then-predict result likely better reflects the true effect size (due to a better signal-to-noise ratio), while the predict-then-average result better demonstrates the consistency of the effect across subjects. These results demonstrate the plausibility of activity flow as a large-scale linking mechanism between resting-state FC and activations across a variety of distinct cognitive tasks. Further, these results suggest a strong role for large-scale distributed (rather than primarily local) processing in the human brain, establishing relevance of resting-state FC to understanding cognitive task activations.

**Figure 2.**
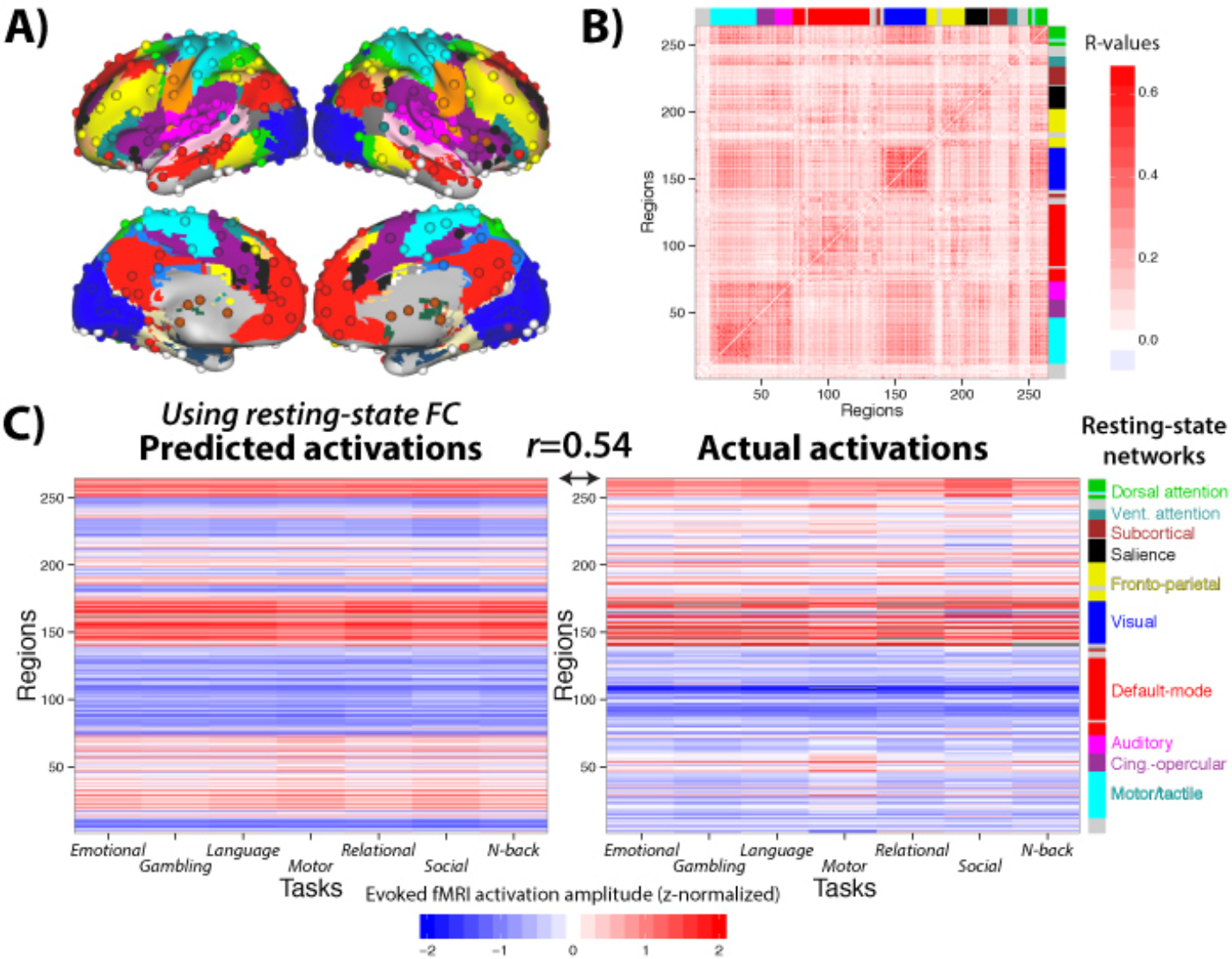
**Activity flow mapping predicts cognitive task activations with empirical fMRI data**. A)We used a standard set of functionally defined regions of interest with associated resting-state FC network assignments^29^. **B**) The across-subject average resting-state FC matrix (Pearson correlations) among the 264 regions shown in panel A. **C**) Resting-state FC was used to predict activation patterns across the 7 tasks (mean activity amplitude of each region for each task). The high correspondence between predicted (left) and actual (right) activation patterns (mean Pearson correlation of r=0.54) suggests resting-state FC shapes activity flow in task contexts.

We next conducted a permutation test to determine whether the observed prediction accuracy was dependent on the particular organization of the FC network architecture. We randomly permuted FC patterns across regions (10,000 permutations; see Methods). We found that the original result was highly dependent on each region’s particular FC pattern. Out of the 10,000 permutations, the highest Pearson correlation r-value between predicted and actual activity was *r*=0.04234. This indicates that the non-parametric permutation test p-value for the original result (*r*=0.54) was ρ<0.0001. **Figure 3A** depicts prediction basedon an example permutation, while **Figure 3B** visually illustrates the null distribution created for the permutation test.

**Figure 3.**
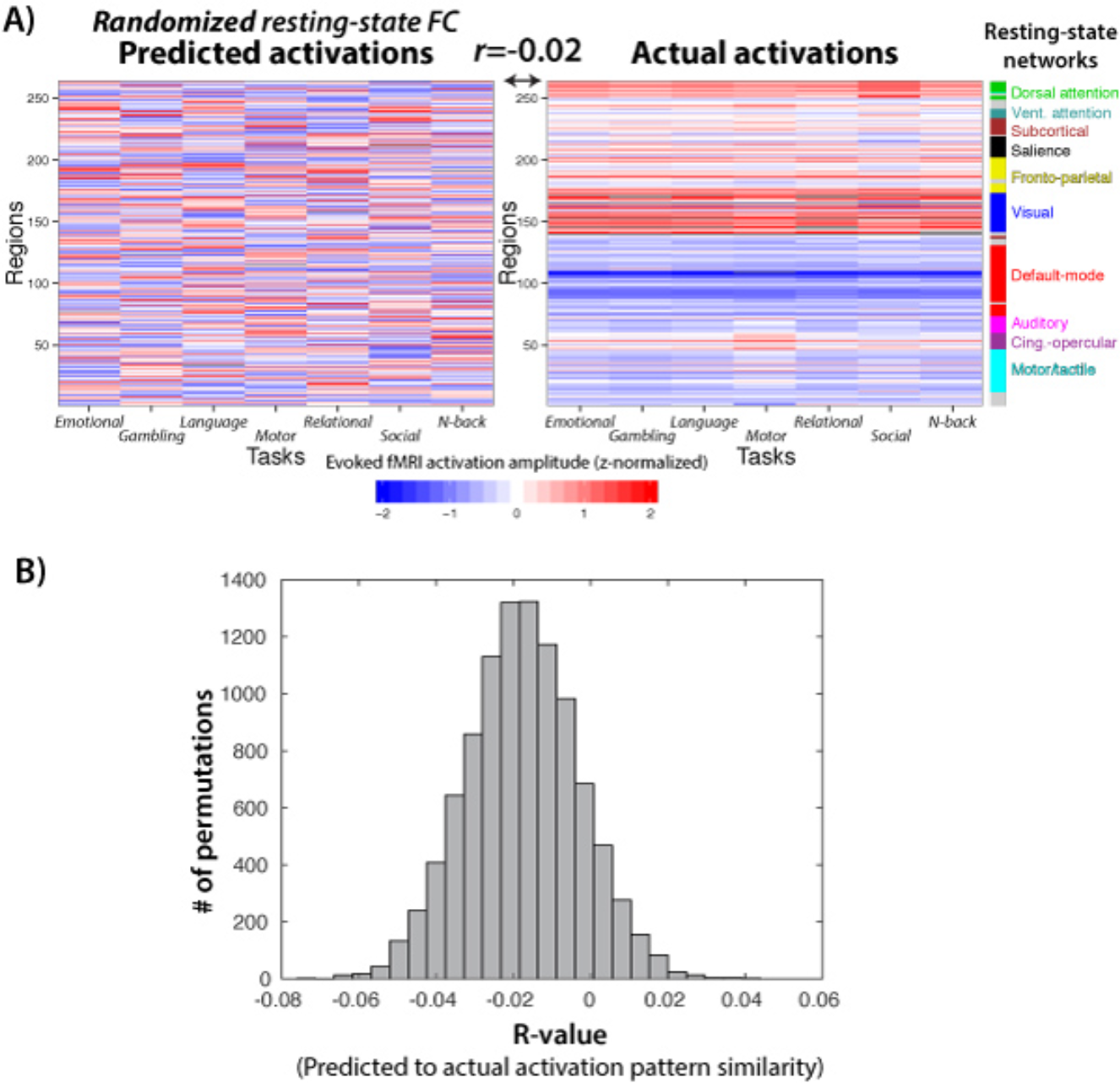
**Activity flow-based predictions depend upon an accurate FC architecture**.We randomized which region’s FC was used for each region’s prediction. **A**)An example of what happens to the activation prediction matrix when FC is randomized. **B**) The distribution of Pearson correlation *r*‐values between predicted and actual activity patterns over 10,000 permutations of FC. The highest *r*‐value was 0.04234.

### Voxelwise activity flow with empirical fMRI data

We next sought to model activity flow between the smallest brain volumes available to us: 2 mm cubic voxels. We performed this analysis to gain a more general assessment of the accuracy of the activity flow modeling approach (e.g., without assuming a set of a priori defined brain regions). Note that we excluded FC with all voxels within the same region (and voxels within 10 mm) of the to-be-predicted voxel to reduce the chance of spatial autocorrelations^30^ contributing to prediction accuracies (see Methods).

We found that whole-brain voxelwise activation patterns were predicted well above chance: average *r*=0.50, t(99)=56.49, ρ<0.00001. This was true for each of the seven tasks individually: *r*=0.50 (emotional), *r*=0.50 (gambling), *r*=0.50 (language), *r*=0.48 (motor), *r*=0.51 (relational), *r*=0.51 (social), *r*=0.48 (N-back). The actual and predicted activation patterns for a single task are depicted in **Figure 4**, visually illustrating the correspondence evidenced by the quantitative results above.

**Figure 4.**
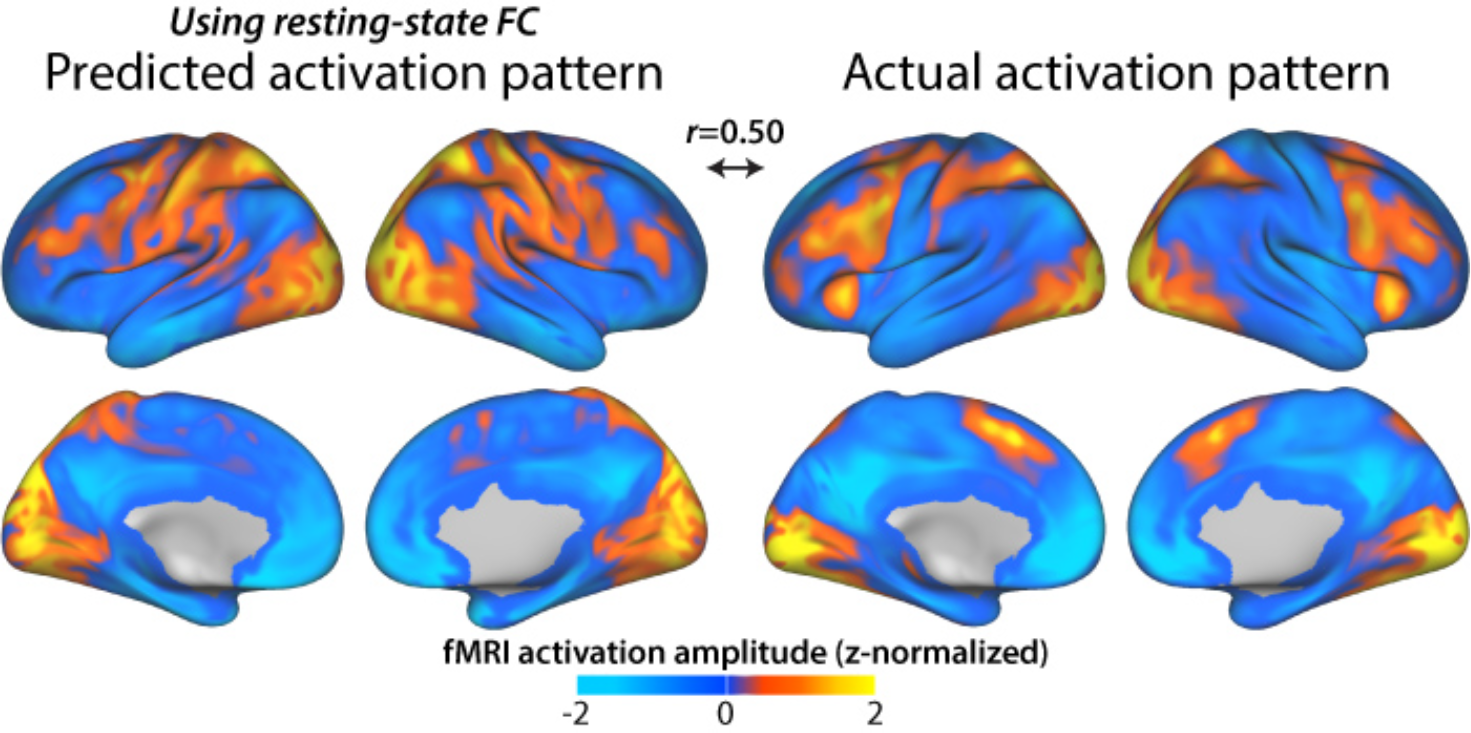
**Predicting voxelwise activation patterns for the N-back working memory task, using resting-state FC.** We applied the activity flow modeling approach in a voxelwise manner, again finding above-chance prediction accuracy across the 7 tasks. Note that voxels within the same region and voxels within 10 mm of the to-be-predicted voxel were excluded from prediction calculations to reduce the influence of spatial autocorrelations (see Methods).

### Improving activity flow mapping predictions using multiple regression

We used Pearson correlations to this point due to their prominent role in the resting-state FC fMRI literature. However, multiple regression is a standard measure for making predictions of a single variable based on many other variables-the goal of the activity flow mapping approach. We therefore adapted the activity flow mapping approach to use multiple regression in place of Pearson correlation. This involved calculating resting-state FC using a standard linear regression model (i.e., a general linear model) for each region, with all other regions as predictor variables. Each regression coefficient in the resulting FC matrix represents how much a given source region’sactivity must be scaled (statistically controlling for all other source regions) to match the activity amplitude of a given target region during resting state.

Using this new FC matrix substantially improved activity flow mapping predictions (**Fig. 5**): average *r*=0.87, t(99)=81.44, ρ<0.00001. The *r*-values were similar for each of the seven tasks individually: *r*=0.87 (emotional), *r*=0.87 (gambling), *r*=0.87 (language), *r*=0.83 (motor), *r*=0.88 (relational), *r*=0.87 (social), *r*=0.87 (N-back). These results demonstrate the utility of using multiple regression rather than Pearson correlations in the context of activity flow mapping. Further, the high correlations obtained further support the possibility that activity flow over intrinsic networks (as estimated by resting-state FC) may strongly shape cognitive task activations.

**Figure 5.**
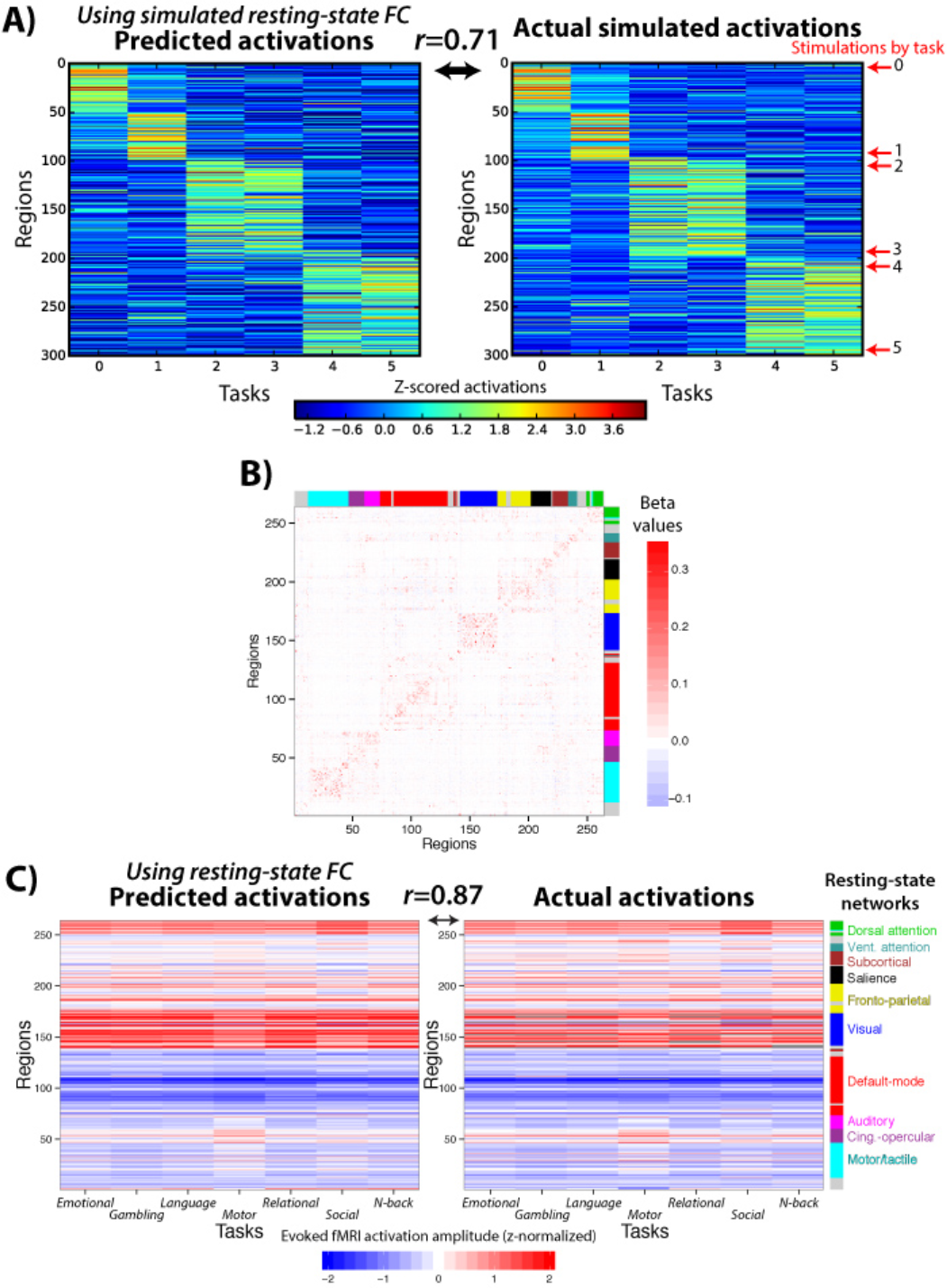
**Using multiple regression to estimate resting-state FC increases prediction accuracy. A**) We applied standard multiple linear regression to estimate each region’s FC in the same simulated data shown in Figure 1. This increased prediction accuracy from *r*=0.56 (with Pearson correlation FC) to *r*=0.71 in this example. **B**) The multiple regression FC matrix from the real resting-state fMRI data. The across-subject average regression coefficient matrix is shown.Some community structure was apparent, despite the increased sparseness relative to when Pearson correlation was used (Fig. 2B). C) Prediction accuracy was also increased with real fMRI data: from an average of *r*=0.54 (using Pearson correlation FC) to an average of *r*=0.87 (with multiple regression FC).

Activity flow mapping predictions were also improved for the voxelwise analysis(**Fig. 6**):*r*= *r*=0.71, t(99)=66.62, ρ<0.00001. The *r*-values were similar for each of the seven tasks individually: *r*=0.71 (emotional), *r*=0.72 (gambling), *r*=0.72 (language), *r*=0.68 (motor), *r*=0.73 (relational), *r*=0.71 (social), *r*=0.70 (N-back). Note that, unlike the regionwise analysis above, it was statistically impossible to include all predictors (here, voxels) for all to-be-predicted voxels. This is due to multiple regression requiring more data points than predictors.We used principal components regression to get around this limitation (see Methods). However, because not all resting-state FC variance was included in the predictions these may be under-estimates of the voxelwise predictions possible with more data. Overall, these results further demonstrate the utility of using multiple regression rather than Pearson correlations in the context of activity flow mapping.

**Figure 6.**
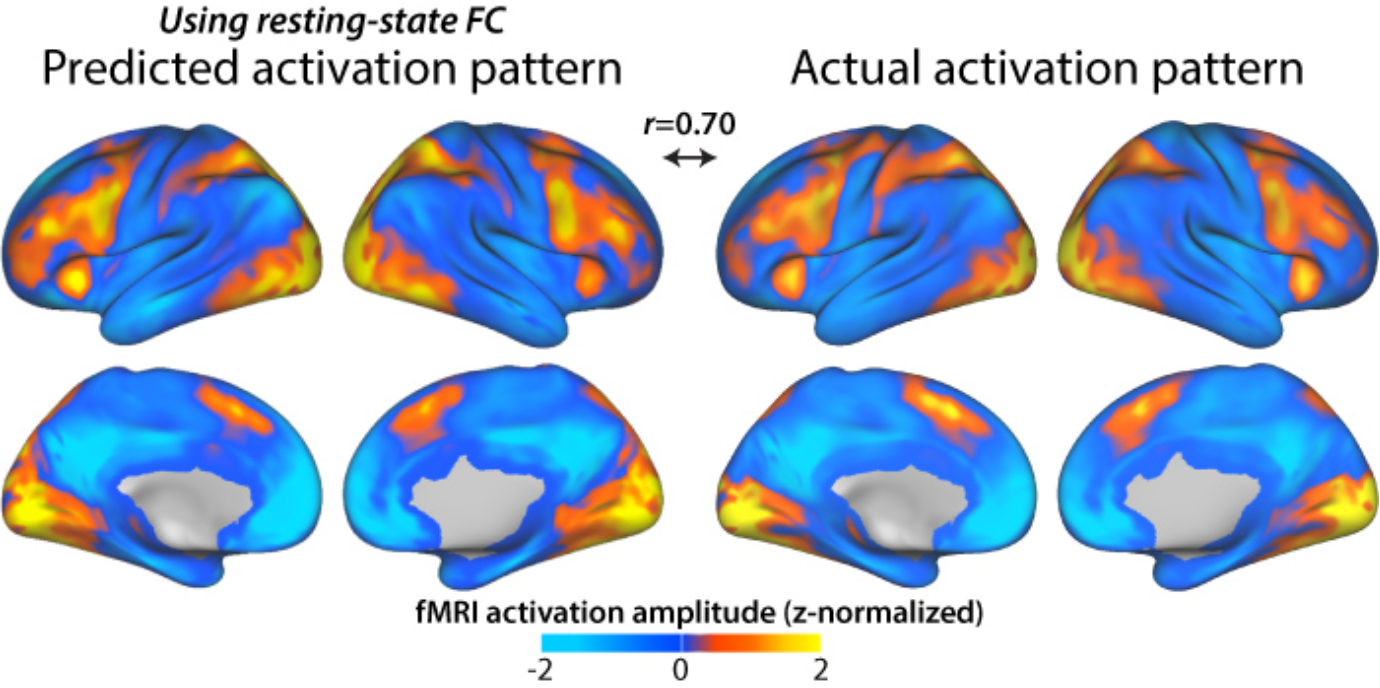
**Predicting voxelwise activation patterns for the N-back working memory task, using multiple regression-based resting-state FC.** We used multiple regression-based resting-state FC with the voxelwise activity flow modeling approach, again finding an increase in prediction accuracy (relative to the Pearson correlation-based FC results) across the 7 tasks. Note that voxels within the same region and voxels within 10 mm of the to-be-predicted voxel were excluded from prediction calculations to reduce the influence of spatial autocorrelations (see Methods). Also, due to fewer time points than predictors, only a subset of the data could be used to compute voxelwise multiple regression FC (see Methods).

### Isolating and predicting task-specific activations

The high similarity among the predicted activation patterns across tasks (see **Fig. 2C**) suggested a potential inability of the activity flow mapping approach for predicting task-specific activations (e.g., motor network activations during the motor task). Similarity was high between tasks in the actual activation patterns here (see **Fig. 2C**), consistent with previous meta-analyses^31  –34^, suggesting the existence of a“task-general” activation pattern. We therefore conceptualized a given task activation pattern as being composed of a task-general pattern and a task-specific pattern (**Fig. 7A**). We used this framework to isolate task-specific activations (e.g., motor network activations during the motor task), which we then used with activity flow mapping. The predicted activation patterns were again well above chance on average: *r*=0.69, t(99)=46.18, ρ<0.00001. The r-values were similar for each of the seven tasks individually: *r*=0.61 (emotional), *r*=0.72 (gambling), *r*=0.64 (language), *r*=0.73 (motor), *r*=0.72 (relational), *r*=0.75 (social), *r*=0.64 (N-back). The average correlation was higher when computed after averaging the FC and activation patterns across subjects (average *r*=0.88; 77% of variance), likely due to improved signal-to-noise from aggregating more data prior to the comparisons (**Fig. 7B**). This was true of each of the seven tasks individually (rvalues, in the same order as above): 0.81, 0.91, 0.86, 0.89, 0.92, 0.89, and 0.84. These values were calculated using multiple regression, given the higher accuracies that were obtained using this approach, though Pearson correlations also provided strong predictions (see Supplementary Information). These results indicate that activity flow over intrinsic networks also shaped observed task-specific cognitive task activations.

**Figure 7.**
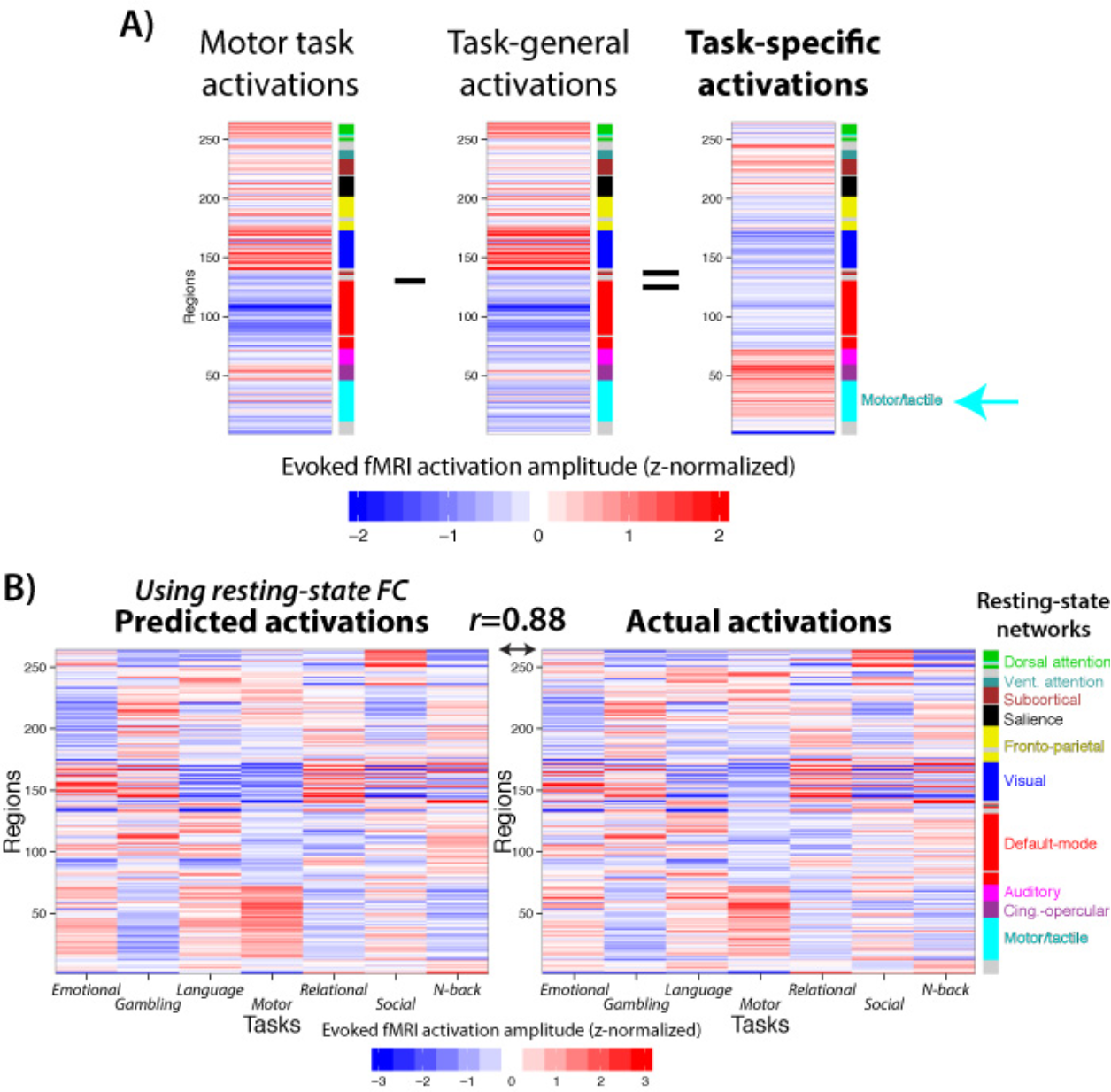
**Isolating and predicting task-specific activations**. **A**) We illustrate a framework involving the decomposition of task activation patterns with the motor task as an example. Whole-brain task activation patterns are shown to be composed of a task-general activation pattern common across tasks (the first principal component of the other 6 tasks) and a task-specific activation pattern (the task activation vector with the task-general activation vector regressed out). Note the task-specific increase in the motor/tactile network (cyan arrow) consistent with the motor task. **B**) Activity flow mapping accurately predicts task-specific brain region activations. This suggests intrinsic networks shape cognitive task-specific activations. Note that multiple regression was used to estimate resting-state FC to improve prediction accuracies, though predictions were also high when using Pearson correlations.

### Predicting individual differences in cognitive task activations

We next tested whether activity flow mapping can be used to predict individual differences in cognitive task activations based on individual differences in resting-state FC. A recent study was able to do this using an abstract statistical model trained to directly associate (within small patches of cortex) resting-state FC values to cognitive task activations ^9^. We postulated that if activity flow is a large-scale mechanism linking resting-state FC to cognitive task activations then activity flow mapping would also produce above-chance prediction of held-out individual subjects. Importantly, unlike the previous study, activity flow mapping does not involve training of an abstract statistical model associating resting-state FC with task activations, potentially demonstrating a more direct relationship between resting-state FC and cognitive task activations.

In addition to holding out each region one-at-a-time, for this analysis we also held out activations from each subject one-at-a-time. This allowed us to use the held-outindividual’s resting-state FC-in combination with other subjects’ mean task activations-to predict the held-out individual’s cognitive task activations (see Methods). The predicted activation patterns were again well above chance on average: *r*=0.76, t(99)=65.92, ρ<0.00001. The r-values were similar for each of the seven tasks individually: *r*=0.77 (emotional), *r*=0.76 (gambling), *r*=0.77 (language), *r*=0.71 (motor), *r*=0.79 (relational), *r*=0.76 (social), *r*=0.76 (N-back). Note that these results were based on multiple regression resting-state FC, though predictions were also accurate when using Pearson correlations (see Supplementary Information). These results demonstrate that resting-state FC describes individualized routes of activity flow, which shape individual differences in cognitive task activations.

It is possible activation predictions in the held-out individuals were above chance due to the general similarity of activations across subjects, rather than due to prediction of individual differences. Consistent with this, across-subject cognitive task activation pattern similarity was *r*=0.63 on average. We therefore isolated individual differences in predicted and actual activation patterns (see Supplementary Information). This revealed that individual differences in cognitive task activations were predicted well above chance on average: *r*=0.39, t(99)=37.76, ρ<0.00001. This was true for all seven tasks separately (see Supplementary Information). These results suggest that although subjects have similar resting-state FC (*r*=0.44 with Pearson correlation FC and *r*=0.40 with multiple regression FC on average) and cognitive task activation patterns (*r*=0.63 on average), resting-state FC nonetheless describes individualized routes of activity flow that shape individual differences in cognitive task activations. See Supplementary Information for additional results also consistent with this possibility.

## DISCUSSION

Recent studies have shown that there is a strong statistical relationship between resting-state FC and cognitive task activations^8, 35^. This was shown using meta-analytic data from thousands of fMRI experiments^8^ and in individual subjects performing specific tasks^9^. However, it has remained unclear how or why this relationship exists. Understanding this relationship in a more mechanistic manner would provide critical insight into the relevance of resting-state FC for cognitive task activations. This would also provide insight into the factors that shape cognitive task activations-a central goal of cognitive neuroscience. Based on our recent work showing that resting-state FC patterns are present during task performance^10^, we expected that resting-state FC may describe the routes of activity flow even during task performance. Building on this, we tested the possibility that activity flow is a linking (large-scale) mechanism between resting-state FC and cognitive task activations, potentially explaining the statistical relationship previously observed between these two constructs.

We found using empirical fMRI data that activity flow across resting-state FC networks could predict cognitive task activations (**Fig. 2**). This was true when holding out each brain region (or voxel), but also when holding out each individual. This demonstrated that individual differences in intrinsic network activity flow can help explain individual differences in cognitive task activations. This may have application in the future for predicting and understanding cognitive task activations in patients who cannot perform a given task (e.g., due to lack of consciousness or cognitive disability) or who perform the task poorly. This addresses a key issue in the study of cognitive disability: We wish to investigate patients with cognitive disabilities using the tasks they have difficulty with, but by definition they will be performing those tasks differently than healthy control subjects.This leads to “performance confounds”, in which any observed change in cognitive task activations could be either a cause or a consequence of the disrupted cognitive task performance. Use of activity flow mapping (and related approaches^9, 14, 15^) may allow us to get around this confound,since we can now understand individual differences in cognitive task activations in terms of connectivity variables estimated independently of task performance.

Several recent studies also sought to identify the relationship between individual subject connectivity and cognitive task (as well as brain stimulation-based^36^) activations^9, 14, 15^. These studies found that functional and structural connectivity patterns-when combined with a statistical model fit to separate data-could be used to predict individual differences in cognitive task activations. These studies provided further evidence that there is a relationship between large-scale connectivity and cognitive activations. Unlike these studies, we utilized a phenomenological construct-activity flow-to link connectivity and cognitive task activity without use of a statistical model trained to relate connectivity to activations. Instead, the relationship between connectivity and cognitive task activations emerged without explicit training of a model to relate these measures. This allowed us to infer a more direct relationship between connectivity and task activations, and further to link this relationship to a potential underlying large-scale mechanism (activity flow, likely shaped by aggregate synaptic connectivity strengths; **Fig. 1D**). Linking to mechanistic constructs (large-scale or otherwise) is important for theoretical advances in neuroscience. In this case, linking to activity flow supports an explanation for the statistical relationships observed in these previous studies. It will be important for future studies to build on these findings with manipulations of FCand activity flow to make more causal inferences about these constructs. Further, it will be important to investigate the relationship between these constructs using more direct measures of neural activity such as multi-unit recording or magnetoencephalography, given that (while strong^37, 38^) the link between neural activity and fMRI BOLD is indirect.

We began by using a simple FC measure (Pearson correlation) to model activity flow.We did this primarily to make minimal assumptions regarding the true nature of brain interactions, and due to widespread use of Pearson correlations for FC estimation in the literature^39, 40^. It is noteworthy that we observed such high accuracy in our predictions (over 50% of variance explained; **Fig. 4**) despite using FC estimates that lack information about both the direction of influence and whether an influence between nodes is indirect (i.e., effective connectivity)^41^. We found that when weused multiple linear regression as an FC measure activity flow mapping accuracies increased (**Fig. 5**). Unlike Pearson correlation this measure takes directionality into account, and also isolates unique influences between regions. Just as we found that using multiple regression FC increased the activation pattern prediction accuracy, so we expect that adding additional (or more accurate) information-perhaps using more sophisticated effective connectivity methods^42 –44^-will improve prediction accuracy further. This would provide evidence for the importance of these factors in shaping cognitive task activations. More generally, this illustrates a benefit of the activity flow framework: the accuracy of predicted activation patterns can be evidence for the veracity of any connectivity properties of interest.

It is important to consider our approach in the context of other modeling frameworks. The activity flow approach is analogous to a model of task activation at time t+1 based on the product of the activation at time t and the connectivity matrix.
As such, this is a single-time step prediction based on the simple linear model utilized in previous studies^25, 45, 46^, with one important change: that the connectivity matrix being utilized is based on FC rather than structural connectivity. Our results may help to explain the observed correlations between cognitive task activation profiles and resting-state FC profiles with fMRI^8^, though they may not explain the lack of correlations between activity and connectivity observed with some electrophysiological neuroimaging approaches^47^. It will be interesting in future to extend our model to include considerations of nonlinear dynamics, such as those implemented in the Virtual Brain project^27, 48^.

To conclude, it is well established that there is a strong statistical relationship between resting-state FC and cognitive task activations^8^, yet the reason for this relationship has remained unclear. We provided evidence for a large-scale mechanism involving activity flow over intrinsic networks (described by resting-state FC) shaping cognitive task activations. This suggests that observed cognitive task activations should not be interpreted simply in terms of localized processing, but should also consider distributed processing in the form of activity flow across intrinsic networks. Further, these results suggest strong relevance of resting-state FC for the task activations that produce cognition. We expect that these insights and the activity flow mapping procedure introduced here will facilitate future investigation into the functional relevance of resting-state FC, the factors that influence cognitive task activations, and the balance of large-scale distributed versus localized processing in the human brain.

## METHODS

**Activity flow mapping**. We developed a method to quantify the relationship between FC and task activation patterns (**Fig. 1A**). This involved estimating net input to each target region by multiplying each other brain region’s task-related activation amplitude (analogous to the amount of neural activity) by its FC with the target region (analogous to aggregate synaptic strength): 
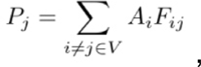
where *P_j_* is the predicted mean activation for region *j* in a given task,*A_i_* is the actual mean activation for region *i* in a given task (a beta value estimated using a general linear model), *i* indexes all brain regions (vector *V)* with the exception of region *j*, and *F_ij_* is the FC estimate between region *i* and region *j* (the Fisher z-transformed Pearson correlation or multiple regression estimate of the regions’ time series). This algorithm results in a vector predicting the pattern of mean activations across regions for a given task. Note that when FC is used rather than effective connectivity this approach estimates total bidirectional (and/or indirect) activity flow. Activation amplitudes were z-normalized for each task separately via subtracting each activation amplitude by the across-region mean and dividing by the across-region standard deviation. This facilitated a focus on the activation patterns (rather than absolute activation levels) across tasks. Prediction accuracy was assessed using Pearson correlation between the predicted activation values and the actual activation values (i.e., the actual activations for each region and task). This was done for each task separately and (unless noted otherwise) each subject separately. Each correlation value was Fisher’s z-transformed prior to averaging, then converted back to a Pearson correlation for reporting purposes. Statistical significance tests were conducted using t-tests (twosided; paired by subject) of Fisher’s z-transformed Pearson correlations, facilitating the ability to infer generalization of results across subjects (rather than just on across-subject mean patterns). The group distributions of these Fisher’s z-transformed Pearson correlations were approximately normally distributed. Whenp-values were computed based on non-normally distributed data we also reported a p-value based on the Spearman’s rank correlation.

**Computational modeling**. We used a simple computational model of large-scale neural interaction to help validate key aspects of activity flow mapping. We sought as simple a computational model as possible to reduce the number of biophysical assumptions and improve the likely generality of our results.

The model consists of 300 abstract units, each representing a brain region. The units interact via a standard spiking rate code passed via predefined structural (and synaptic) connectivity^49^. Activity at agiven node is determined using a standard sigmoid function on the mean of the input activities. Note that the sigmoid function introduces a non-linearity to the interactions among units that is similar to aggregate non-linearity from neuronal action potentials^50^. Specifically, the model used the following equation to determine activity in a given unit at a given time step:

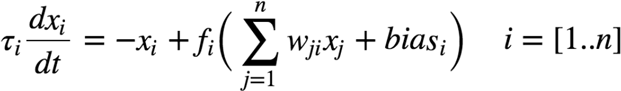
where *w_ij_* refers to the synaptic weight from neuron *i* to *j*, *x_j_* refers to the activity level at region *j*. bias, is the bias of neuron *i,* but for this model this is set to 0. *τ_i_* is the time constant for region *i*, and is set to 1 time step for all regions.

The model’s network connectivity was constructed by first defining a random set of structural connections (15% density), then creating 3 graph communities/sub-networks by randomly connecting each node to 10 other nodes within the same community.Structural connections were defined as non-zero connection weights (all set to the same value of 1.0), while synaptic connections were modifications on the initial connection weight. Normally distributed random synaptic weights were added to all structural connections, scaled to be quite small (mean of 0 and standard deviation of 0.001). Finally, synaptic weights were used to split the first structural connectivity community into two “functional” communities. Specifically, the synaptic weights were increased (multiplied by 1.5) within the first half of the first graph community, while synaptic weights for the second half of that community were also increased (multiplied by 1.5). Also, synaptic weights between these communities were reduced (multiplied by 0.5). These modifications were designed to test the impact of synaptic weights on simulated activity flow.

Spontaneous activity for each node was added as normally distributed random values every time step (100 ms). An autocorrelation factor of 0.10 was used to maintain some activity across multiple time steps. 20,000 time steps were simulated using purely spontaneous activity (resting state data) and, separately, using spontaneous with task-evoked activity (task data). Task-evoked activity was implemented as increased activity (normally distributed random values centered at 1 with a standard deviation of 0.5) added linearly to ongoing spontaneous activity. Activity consisted of 3 blocks of 2,000 time points each, each separated by 3,000 time points. Each task was simulated by adding task-evoked activity to six separate groups of five regions simultaneously (2 per structural graph community). FMRI data collection was simulated by convolving the simulated time series with the SPM canonical hemodynamic response function, then downsampling to a standard TR of 2 seconds. All analyses of the simulated fMRI data were identical to the analyses conducted on the empirical fMRI data.

We defined a global coupling parameter as a scalar multiplier on all synaptic strengths, and a local processing parameter as a scalar multiplier of all self-connection strengths. Self-connections increase the influence of a region’s activity on itself in the next time step, separating variance in its activity from the activity of other regions. For the parameter sweep (**Fig. 1B**) we used 20 global coupling parameters (from 0-5, using 0.25 increments) and 20 local processing parameters (from 0-100, using 5.0 increments), each averaged across 10 “subjects” (separate iterations with random initial structural/synaptic connectivity matrices). This totaled 4,000 simulations. Modeling was carried out using Python (version 2.7).

**Data collection**. Data were collected as part of the Washington University-Minnesota Consortium Human Connectome Project^51^. Human participants were recruited from Washington University (St. Louis, MO) and the surrounding area. All participants gave informed consent consistent with policies approved by the Washington University Institutional Review Board. The data used were from the “500 Subjects” HCP release. The “100 Unrelated Subjects” (N=100) subset of this dataset was used, given that a subset of unrelated individuals is more appropriate for statistical analyses intended to represent the general population. Based on our primary statistical tests (paired t-tests,alpha=0.05) and assuming a moderate Cohen’s d effect size of 0.5, N=100 provides 99.86% power^52^ (higher than the standard criterion of 80%). The average age of the participants was 29 years (ranging from 22 to 36), with 54% females. Whole-brain echo-planar imaging acquisitions were acquired with a 32 channel head coil on a modified 3T Siemens Skyra with TR=720 ms, TE=33.1 ms, flip angle=52°, BW;=2290; Hz/Px, in-plane FOV=208×180 mm, 72 slices, 2.0 mm isotropic voxels, with a multi-band acceleration factor of 8^53^. Data were collected over two days. On each day 28 minutes of rest (eyes open with fixation) fMRI data across two runs were collected (56 minutes total), followed by 30 minutes of task fMRI data collection (60 minutes total). Each of the 7 tasks was completed over two consecutive fMRI runs. Resting-state data collection details for this dataset can be found elsewhere^54^, as can task data details^28^.

**Code availability.** The code used for activity flow mapping and computational modeling will be made available on our lab website (http://www.colelab.org) and GitHub (https://github.com/ColeLab/).

**Data availability**. All data are available at http://humanconnectome.org/.

**Task paradigms**. The dataset was collected as part of the Human Connectome Project, and included rest and a set of seven tasks^28^. These tasks included seven distinct domains: emotion, reward learning, language,motor, relational reasoning, social cognition, and working memory. Briefly, the emotion task involved matching fearful or angry faces to a target face. The reward learning task involved a gambling task with monetary rewards and losses. The language task involved auditory stimuli consisting of narrative stories and math problems, along with questions to be answered regarding the prior auditory stimuli. The motor task involved movement of the hands, the tongue, and the feet. The relational reasoning task involved higher-order cognitive reasoning regarding relations among features of presented shape stimuli. The social cognition (theory of mind) task used short video clips of moving shapes that interacting in some way or moving randomly, with subjects making decisions about whether the shapes had social interactions. The working memory task involved a visual n-back task, in which subjects indicate a match of the current image to either aconstant target image or two images previous.

**Data preprocessing**. Preprocessing consisted of standard resting-state functional connectivity preprocessing (typically performed with resting-state data), with several modifications given that analyses were also performed on task data. Resting-state and task data were preprocessed identically in order to facilitate comparisons between them.

Spatial normalization to a standard template, motion correction, and intensity normalization were already implemented as part of the Human Connectome Project in a minimally processed version of the dataset described elsewhere^55^. With the volume(rather than the surface) version of the minimally preprocessed data, we used AFNI^56^ to additionally remove nuisance time series (motion, ventricle, and white matter signals, along with their derivatives) using linear regression, remove the linear trend for each run, and spatially smooth the data. The data were smoothed using a non-Gaussian filter (nearest neighbor averaging) at 4 mm to reduce the chance of introducing circularity in the activity flow mapping procedure (see below). Unlike some standard resting-state FC preprocessing pipelines, whole brain signal was not included as a nuisance covariate (given current controversy over this procedure^57^), and a low-pass temporal filter was not applied. We did not apply a low-pass temporal filter given the likely presence of task signals at higher frequencies than the relatively slow resting-state fluctuations, and the desire to preprocess the rest and task data similarly. Note that activity flow mapping results were similar after whole brain signal regression. Freesurfer^58^ was used to identify ventricle, white matter, and gray matter anatomical structures for each participant.

For the main analyses, data were sampled from a set of 264 brain regions (rather than individual voxels) in order to make inferences at the region and systems level (**Fig. 2A**). We used an independently identified set of putative functional brain regions^29^ rather than anatomically defined sets of regions in order to reduce the chance of combining signal from multiple functional areas^59^. These brain regions were identified using a combination of resting-state FC parcellation^60^ and task-based neuroimaging meta-analysis^29^. Data were summarized for each region by averaging signal in all voxels falling inside each region. Analyses were carried out with MATLAB 2014b (Mathworks) and R 3.1.2 (The R Foundation for Statistical Computing).

**FC estimation**. The initial analyses estimated FC using Pearson correlations between time series (averaging across voxels within each region) from all pairs of brain regions. The same procedure was used for the voxelwise analyses, but between voxels rather than regions. All computations used Fisher’s z-transformed values, which were reconverted to r-values for reporting purposes.

We used standard multiple linear regression (the regstats function in MATLAB) as an alternative to Pearson correlation. This involved computing a linear model for each to-be-predicted region separately. Resting-state fMRI time series from all other regions were used as predictors of the to-be-predicted region’s resting-state fMRI timeseries. The resulting betas-which were directional from the predictor regions to the predicted region - were then used as FC estimates in the activity flow mapping algorithm.

**Task activation level estimation**. The activation amplitudes were estimated using a standard general linear model. The SPM canonical hemodynamic response function was used for general linear model estimation, given that all tasks involved block designs.

**Activity flow model permutation testing**. We used permutation testing to help validate the activity flow mapping approach, and provide an additional means of inferring statistical significance. The permutation test was constructed so as to facilitate a conservative statistical inference, wherein only the hypothesized essential aspect of the analysis was permuted. This involved keeping all aspects of the analysis
the same except for random permuting (without replacement) which region’s FC was used on each iteration. In other words, the entire set of FC strengths for the to-be-predicted region was swapped with the entire set of FC strengths for another region chosen uniformly at random from the set of all regions. This permutation process was run 10,000 times (with resting-state FC), resulting in a null distribution of r-values centered just below 0 and with a maximum value of 0.04234. Thus a prediction accuracy r-value above 0.04234 would be significantly at the level of ρ<0.0001.

**Voxelwise activity flow mapping**. We made relatively minimal changes tothe regional activity flow modeling when applying it in a voxelwise manner. First, we excluded all voxels within 10 mm of the to-be-predicted voxel to reduce the chance of spatial autocorrelations contributing to prediction accuracies^30^. Second, we excluded all voxels within the same functional region (defined as local voxels with similar resting-state FC patterns) as the to-be-predicted voxel in order to reduce the influence of potentially trivial within-region activity flow upon prediction accuracies. The recently developed Gordon cortical area parcellation^61^ was used because (unlike the Power brain area parcellation used for the other analyses) it included a voxelwise version amenable to our processing pipeline and because its development involved similar principles as the Power brain area parcellation. Finally, the voxelwise activity flow predictions were calculated for each subject independently, and the resulting prediction maps were subsequently averaged across subjects. Results are also reported with predictions compared to actual activation patterns for each subject separately.We used Connectome Workbench software (v1.0) for visualization.

For the multiple regression-based voxelwise activity flow approach, there were many more predictors (voxels) than time points. Thus, unlike the regionwise analyses, this made it impossible to compute FC estimates using all available predictors. Instead we used a standard statistical approach for performing multiple regression with many more predictors than data points: principal components regression^62^. Briefly, this involved extracting the time series for the first 500 principal components, performing the regression on each voxel using those components, then projecting the resulting beta values back into the original voxel space (rather than the principal component space). We used the first 500 components and the first 1200 resting-state fMRI time points for computational tractability.

**Prediction of individualized task activations**. Each subject’s cognitive task activations were held out in a leave-one-subject-out cross-validation approach. The held-out individual’s resting-state FC-along with other subjects’ task activations-were used to predict the held-out individual’s cognitive task activations. Specifically, task activations were averaged across all subjects except the held-out subject, then the activity flow mapping procedure was applied along with the held-out subject’s resting-state FC. This allowed us to quantify the likely role of that individual’s intrinsic connections (as estimated by resting-state FC) in shaping cognitive task activations.

**Task-specific activation patterns**. Task-general activation patterns were defined as the first principal component across task activation patterns. Principalcomponent analysis was used rather than averaging to reduce the chance that any individual task’s activation pattern dominated the task-general pattern. This was computed separately for each subject, and also for each task; the to-be-predicted (or compared) task’s activations were excluded to remove circularity from the calculation. Results were virtually identical if all seven tasks were included in the task-general activation calculation. Task-specific activation patterns were defined as a given task’s activation pattern after regressing out task-general activations (the first principal component across the other six tasks’ activation patterns). The average pairwise similarity among task-specific activation patterns (i.e., after regressing out task-general activations) was r=‐0.1.

## Acknowledgements

We would like to thank Bharat Biswal, Todd Braver, Steven Petersen, and Jonathan Power for helpful conversations during preparation of this manuscript. Data were provided by the Human Connectome Project, WU-Minn Consortium (Principal Investigators: David Van Essen and Kamil Ugurbil; 1U54MH091657) funded by the 16 NIH Institutes and Centers that support the NIH Blueprint for Neuroscience Research; and by the McDonnell Center for Systems Neuroscience at Washington University. M.W.C. was supported by the US National Institutes of Health under award K99-R00 MH096801. D.S.B. acknowledges support from the John D. and Catherine T. MacArthur Foundation, the Alfred P. Sloan Foundation, and the US Army Research Office through contract no. W911NF-14-1-0679. The content is solely the responsibility of the authors and does not necessarily represent the official views of any of the funding agencies.

## SUPPLEMENTARY INFORMATION

### Isolating and predicting task-specific activations using Pearson correlation FC

We also predicted the task-specific activations using Pearson correlation FC estimates, since this better relates results to the existing resting-state FC fMRI literature. The predicted activation patterns were again well above chance on average (Supplemental Fig. 1): *r*=0 .48, t(99)=39.29, ρ<0.00001. The r-values were similar for each of the seven tasks individually: *r*=0.42 (emotional), *r*=0.49 (gambling), *r*=0.46 (language), *r*=0.53 (motor), *r*=0.49 (relational), *r*=0.50 (social), *r*=0.45 (N-back). Thus, the results indicate that activity flow over intrinsic networks (as estimated using resting-state Pearson correlations) also shaped observed task-specific cognitive task activations.

**Figure S1.**
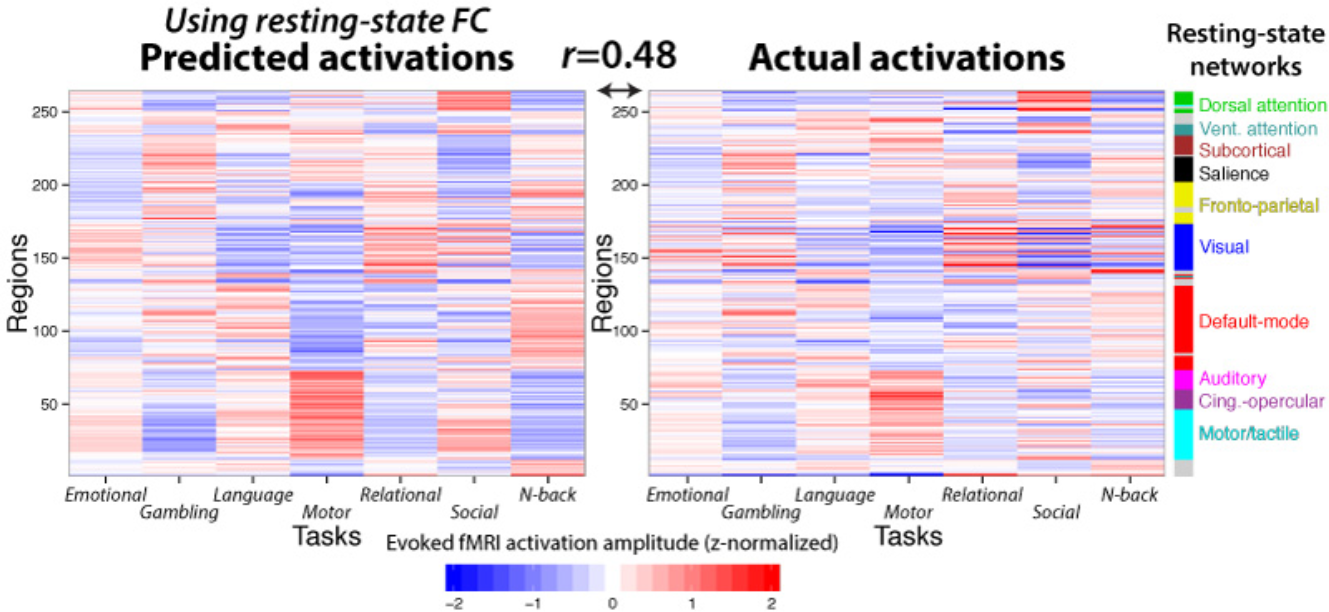
**Isolating and predicting task-specific activations using Pearson correlation FC.** Activity flow mapping using Pearson correlations accurately predicts task-specific brain region activations. This mirrors the result using multiple regression FC (Fig. 7), though with lower accuracy (as expected).

**Predicting individual differences in cognitive task activations using Pearsoncorrelations**We also predicted individual differences in cognitive task activations using Pearson correlation FC estimates, since this better relates results to the existing resting-state FC fMRI literature. The predicted activation patterns were again well above chance on average: *r*=0.48, t(99)=38.01, ρ<0.00001. The r-values were similar for each of the seven tasks individually: *r*=0.50 (emotional), *r*=0.48 (gambling), *r*=0.49 (language), *r*=0.46 (motor), *r*=0.51 (relational), *r*=0.48 (social), *r*=0.48 (N-back). These results demonstrate that resting-state FC (as estimated by Pearson correlation) describes individualized routes of activity flow, which shape individual differences in cognitive task activations.

To more rigorously demonstrate predictions of individual differences, we next assessed whether the predicted activations were more similar for the predicted individuals relative to others. As expected, we found that average predictions dropped to *r*=0.40 when predicting activations of other subjects. This was significantly lower than the original value: t(98)=6.01, ρ<0.00001. Thisresult suggests that although subjects have highly similar resting-state FC and cognitive task activation patterns, resting-state FC nonetheless describes individualized routes of activity flow that shape individual differences in cognitive task activations.

**Isolating individual differences in cognitive task activations prior to prediction comparison** It is possible activation predictions in the held-out individuals were above chance due to the general similarity of activations across subjects, rather than due to prediction of individual differences. Therefore, to more rigorously demonstrate predictions of individual differences, we next assessed whether the predicted activations were more similar to the actual activations for the predicted individuals relative to other individuals. As expected, we found that average prediction accuracy dropped to *r*=0.63 when using any one individual’s resting-state FC to predict activations of other subjects. This was significantly lower than the original value: t(98)=16.67, ρ<0.00001. Note that this value was equal to the amount of between-subject similarity among the actual cognitive activation patterns on average (*r*=0.63), signifying that the prediction accuracy dropped to the baseline level of across-subject cognitive task similarity. These results suggest that although subjects have similar resting-state FC (*r*=0.44 with Pearson correlation FC and *r*=0.40 with multiple regression FC on average) and cognitive task activation patterns (*r*=0.63 on average), resting-state FC nonetheless describes individualized routes of activity flow that shape individual differences in cognitive task activations.

To further support this conclusion, we also removed the subject-general activation pattern for each task prior to assessing prediction performance. This better isolates the subject-specific activation patterns, allowing us to better assess prediction accuracy of these patterns. Subject-general activations were identified as the first principal component across subjects (for a given task). This subject-general pattern was then regressed out of each subject’s activation pattern. This approach was similar to the task-specific pattern isolation approach in a previous analysis. Unlike the task-specific approach, this approach was applied separately for the predicted activation patterns (the same patterns reported in the main individual differences analysis) and the actual activation patterns (for comparison with the predicted patterns). After removing the task-general activation patterns, across-subject activation similarity dropped from *r*=0.63 to *r*=‐0.01. Using these isolated subject-specific activations (with multiple regression FC), we found that the predicted activation patterns were again well above chance on average: r=0.39, t(99)=37.76, ρ<0.00001. The r-values were similar for each of the seven tasks individually: *r*=0.40 (emotional), *r*=0.39 (gambling), *r*=0.41 (language), *r*=0.35 (motor), *r*=0.41 (relational), *r*=0.40 (social), *r*=0.40 (N-back). These results further confirm the ability to use activity flow mapping to predict individual differences in cognitive task activations.

